# Beyond Averaging: Pleiotropic effects of *Dry2.2* expose the evolution of growth bet-hedging in barley

**DOI:** 10.64898/2026.06.22.733768

**Authors:** Ayelet Kurtz-Sohn, Lamis Abdelhakim, Avital Beery, Manas R. Prusty, Klára Panzarová, Eyal Fridman

## Abstract

**Rationale:** Developing climate-resilient crops requires understanding the developmental and physiological mechanisms governing responses to unpredictable environmental stresses. While agricultural selection historically favored phenotypic uniformity, highly volatile conditions can favor risk-spreading “bet-hedging”. However, because standard genetic mapping relies on shifting plot-level averages, loci controlling population-level variance remain cryptic.

**Methods:** We investigated the pleiotropic effects of the *Dry2.2* quantitative trait locus (candidate gene *HvCEN*) using an allelic series of wild barley (*Hordeum vulgare* ssp. *spontaneum*) introgressions housed in distinct cultivated backgrounds, evaluated via high-resolution single-plant physiological phenotyping and mini-plot field trials.

**Results:** Specific wild alleles confer robust developmental canalization under water limitation, maintain harvest traits, stabilize vascular lignification, and drive a uniform senescence escape strategy. Conversely, carriers of cultivated alleles deploy a bet-hedging strategy under stress, more than doubling inter-plant developmental variation (volume, maturity timing). This variance-driven strategy relies on epistatic interactions, rendering *Dry2.2* invisible to traditional mean-centric GWAS plots.

**Conclusion:** Improving crop resilience in volatile climates requires expanding selection focus beyond static, plot-level averages to include the active genetic design of population-level variance strategies.

## Introduction

One emerging question that has long baffled evolutionary biologists, and is becoming increasingly relevant under climate change, is whether plasticity or robustness in core activities is more beneficial for organisms’ fitness at the individual or group level (Alpert and Simms, 2002). Plants, as stationary organisms, have many adaptations to environmental changes, and these responses can be categorized as plasticity (Brooker et al., 2022) or robustness (Lachowiec et al., 2016). Plasticity is defined as the adjustment of a phenotype for the same genotype in response to environmental changes. Robustness, on the other hand, is the ability to maintain stability despite such changes (Waddington, 1942; Lempe et al., 2013; Lachowiec et al., 2016). The interplay of plasticity and robustness at molecular and genetic levels, as well as across different traits, is essential for plant development (Debat and David, 2001). Challenges in understanding evolution, genetics, and the mechanisms of plasticity include defining relevant phenotypes, identifying environmental cues, selecting which molecular layers to examine, and determining whether the unit of selection is the individual or a group (Laitinen and Nikoloski, 2019).

While plasticity is typically viewed as a directed ‘sense-and-switch’ mechanism in response to reliable environmental cues (Alpert and Simms, 2002; Brooker et al., 2022), this uniform strategy can be perilous when environmental fluctuations are highly unpredictable. In such volatile conditions, evolutionary pressures can favor a distinct and subtle manifestation of plasticity known as bet-hedging. Rather than producing a unified phenotypic shift, bet-hedging drives a single genotype to intentionally generate a diverse range of phenotypes within a group. This strategy acts as an evolutionary safeguard by spreading developmental risk; it ensures that at least a fraction of the individuals will survive unpredictable stresses, effectively trading immediate, maximized individual growth for long-term population stability. While often studied in bacteria as stochastic “phenotype-switching” (de Groot *et al., 2023*), in plants, this is most evident in traits such as seed dormancy and germination timing (Abley et al., 2024). For wild species, maintaining a range of individual phenotypes within a population ensures that at least some offspring survive fluctuating conditions, a form of balancing selection that trades immediate maximum growth for long-term population stability.

The broader debate about plasticity is particularly important for global food security. In plants, responses to water limitations often show plasticity, with the organism prioritizing survival over yield. Overall, plasticity is believed to be selected against during domestication. Crossing crop wild relatives with their cultivated counterparts can yield new, potentially beneficial combinations; therefore, wild lines are considered a valuable resource for developing resilient crops (Zamir, 2001; Maurer et al., 2015). Interspecific populations are not only instrumental in capturing and realizing the breeding potential of wild alleles in a relatively cultivated genetic background (Alseekh et al., 2017; Beche et al., 2020), but also in facilitating the identification of a signature of selection related to control of plasticity and robustness. For example, the thermal plasticity of the circadian clock, in which cultivated and landrace barley have lost their period and amplitude responses to higher temperatures, was linked to selected drivers of clock (DOC) loci, which play a key role in mediating these differences (Prusty et al., 2021). Regarding bet-hedging plasticity, it is assumed that it has been subjected to negative selection during domestication, favoring rapid and uniform germination and field establishment, even under sub-optimal conditions (Mitchell et al., 2017).

For a commercially significant crop like barley (*Hordeum vulgare* ssp. *vulgare*), plastic responses may be seen as detrimental, as agricultural success is measured by grain quantity and uniformity (wide adaptation) rather than plant persistence. Cultivated barley has been selected for millennia for traits such as grain yield and day-length insensitivity (Badr et al., 2000; Tao et al., 2022), leading to a narrow genetic pool that lacks the robustness needed to withstand environmental changes.

A genome-wide association study (GWAS) of the multi-parent interspecific HEB-25 population (Maurer et al., 2015) identified a specific allele of the quantitative trait locus (QTL) *Dry2.2* that appears to contribute to robustness in drought-induced, flowering-independent grain number production (Merchuk-Ovnat et al., 2018). This locus is located on the long arm of chromosome 2 and includes several genes involved in flowering. Specifically, HvCEN, an ortholog of the Garden Snapdragon (*Antirrhinum majus*) CENTRORADIALLIS (CEN) (Bradley et al., 1996) and Arabidopsis thaliana’s TFL (Hanano and Goto, 2011). HvCEN has also been linked to flowering and plant architecture (Comadran et al., 2012; Bi et al., 2019; Wang et al., 2020; Göransson et al., 2021). It is a strong candidate gene for the observed yield robustness, as it and its orthologs are involved in drought response (Wang et al., 2020), grain yield (Bi et al., 2019, 20), and inflorescence architecture (Bradley et al., 1996).

Here, we sought to understand plant responses at multiple levels, including in additional wild alleles, to clarify the processes and molecular mechanisms that maintain drought robustness. Surprisingly, these detailed analyses of several nearly isogenic advanced backcross lines, as well as their cultivated recipients, which represent modern cultivars, revealed growth bet hedging under water limitations linked to the cultivated, rather than the wild, allele of *Dry2.2*.

## Materials and Methods

### Plant material and growth conditions

To standardize nomenclature throughout this study, advanced backcross lines are referred to by their recurrent cultivated background followed by their allelic state at the *Dry2.2* locus. For example, homozygous wild introgression lines in the Barke background are denoted as Barke-*Dry*2.2^Hs^, while their sibling lines homozygous for the cultivated allele are denoted as Barke-*Dry*2.2^Hv^. We also distinguish between different wild alleles, e.g., Barke-*Dry*2.2^Hs-HID062^ refers to the introgression of *Dry2.2* from HID062. Lines were advanced to at least the BC_3_S_2_ generation to ensure homozygosity at the target locus, which was confirmed via TaqMan genotyping prior to experimental use. We selected plant lines from the interspecific cytonuclear multi-parent population (CMPP) bred from the B1K collection (Bodenheimer et al., 2025), and from the HEB-25 population bred by Maurer *et al* (2015). Noga and Barke were used for crosses and as controls for the CMPP and HEB lines, respectively. CMPP lines were selected for crossing based on their 50K iSelect SNP genotyping (Bayer et al., 2017), inversion status, and TaqMan genotyping (Fig. 1, Table 1). Homozygous doubled haploids were then backcrossed to Noga, and their offspring were allowed to self-pollinate, yielding fixed progeny for the experiments (Fig. S1).

**Figure 1.**
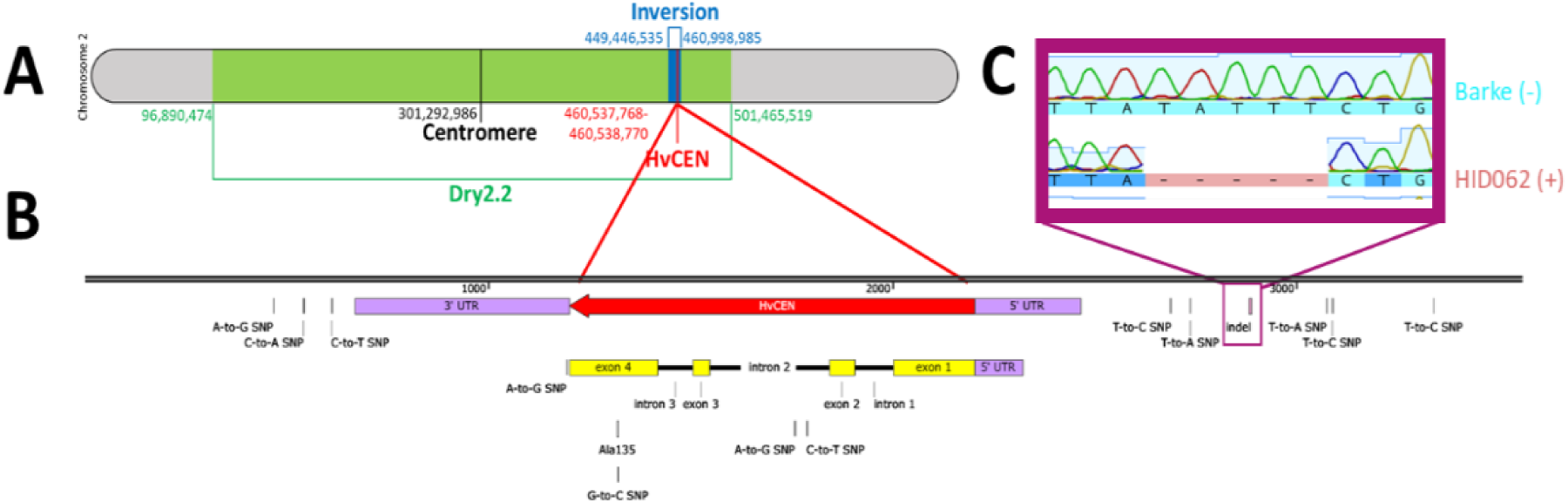
Map of *Dry2.2* and HvCEN. A) *Dry2.2* was identified between markers SCRI_RS_144592 and SCRI_RS_165574 (Merchuk-Ovnat et al., 2018). Within this locus lies an inversion (Jayakodi et al., 2020). B) Sequencing revealed polymorphisms in and around HvCEN, including a non-synonymous SNP in Exon 4 and a deletion in the promoter region. C) An indel was identified in the promoter region of the gene with 5 bp that exists in the cultivated Barke allele but not in the wild HID062 allele named CHAT (*Hv**C**EN* **Ha**kuna **T**ata)

**Table 1:**
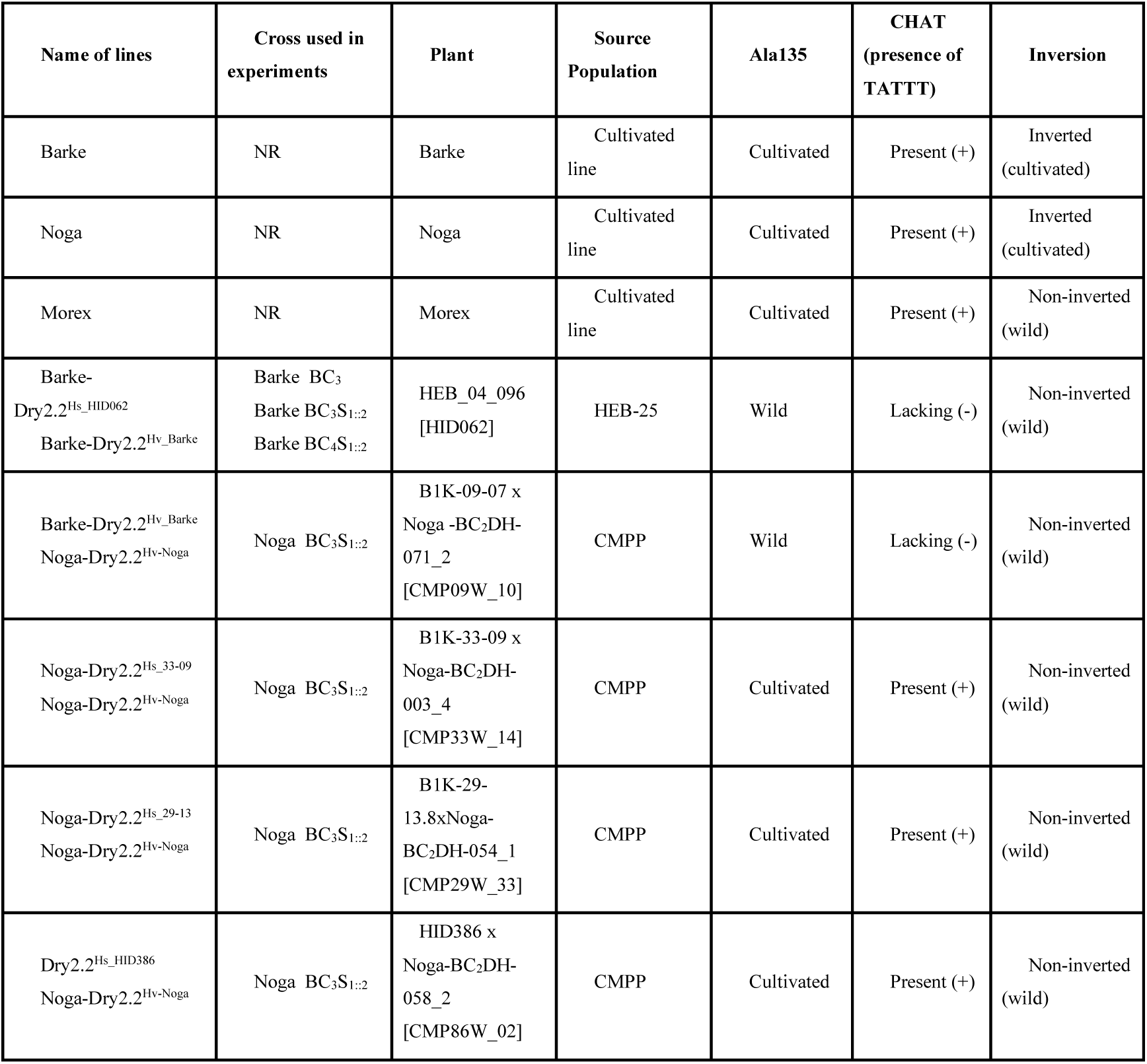
Plant lines used in the study.

Lines selected for use were: *i)* Barke-*Dry*2.2^Hs_HID062^ and Barke-*Dry*2.2^Hv_Barke^ from the 3^rd^ and 4^th^ backcrosses to Barke; *ii)* Noga-*Dry*2.2^Hs_09-07^ and Noga-*Dry*2.2^Hv-Noga^ from BC_3_S_1::2_ of CMPP line CMP09W_10 to Noga, *iii)* Noga-*Dry*2.2^Hs_33-09^ and Noga-*Dry*2.2^Hv-Noga^ from BC_3_S_1::2_ of CMPP line CMP33W_14 to Noga, *iv)* Noga-*Dry*2.2^Hs_29-13^ and Noga-*Dry*2.2^Hv-Noga^ from BC_3_S_1::2_ of CMPP line CMP29W_33 to Noga, and, *v)* Noga-*Dry*2.2^Hs_HID386^ and Noga-*Dry*2.2^Hv-Noga^ from BC_3_S_1::2_ of CMPP line CMP86W_02 to Noga (Table 1).

For both the whole-plant phenotyping and lignin accumulation experiments, seeds were sown in small pots and incubated at 4°C for 2 days to standardize germination. The trays were then transferred to short-day conditions until seedlings reached the 4^th^ leaf stage, after which they were transplanted to troughs in the greenhouse. Plants were grown under natural light conditions for 6 months (December-May, 2021-2 [lignin] 2022-3 [whole plant]) in a Mediterranean climate (Volcani Institute, Rishon-Letziyon, Israel).

Following plant establishment, water limitation was imposed by withholding water, then by providing limited watering. Soil water content was measured using Decagon Devices ProCheck. Water saturation (WW) was defined as VWC (volumetric water content) ≥30, while water limitation (WL) was defined as <15 VWC.

For the time-lapse growth phenotype experiment, after incubation at 4°C for 2 days, plants were transferred to a lit walk-in chamber (FytoScope FS-WI, Photon Systems Instruments (PSI), Drásov, Czech Republic) and grown under short days until the 5^th^ leaf emerged. Plants were then transferred to a 3-L pot filled with 1,850 g of Klasmann Substrate-2:sand (3:1) and grown under long-day (16/8 L/D) conditions for 97 days (September-December 2023 and 2024). Day/Night temperatures were 22°C ± 3°C (mean ± standard deviation)/17 °C ± 2 °C. Day/night relative humidity was 51 ± 8/62 ± 4% for day/night (Tietze et al., 2025).

### Plant Genotyping

For all PCRs, we used Vazyme’s 2 × Rapid Taq Master Mix (Vazyme, Nanjing, PRC). Barke, Noga, and HEB-04-96 were sequenced for ∼3500 bp around *HvCEN* in 5 intervals. The primer pairs used are described in Supplementary Table S1. Inversion was identified by PCR using a triplicate set of three primers: one positioned on one edge of the inversion and one that flanked it on either side (Table S1).

High-resolution melt (HRM) using Quantabio’s AccuMelt^TM^ HRM SuperMix (Quantabio, Beverly, Massachusetts, USA) on the Qiagen Rotor-Gene identified the presence/absence of Indels. Controls for these experiments included Barke and HEB-04-096.

Taqman analysis was performed to identify a SNP in exon 4 on the StepOne Plus, using the TaqPath^TM^ ProAmp^TM^ Master Mix (Thermo Fisher, Waltham, Massachusetts, USA).

### Data collection and analysis

For whole-plant phenotyping, spikes were collected by hand and counted. Next, all spikes were weighed together, and single spike weight was calculated by dividing the total spike weight by their number. Grains were cleaned using an RC52 rice-milling machine (Yamamoto Electric, Fukushima, Japan). The grains were separated from the chaff using a hairdryer on a low setting. The cleaned seeds were then weighed. Grains were counted using the Count Things app (Professional Counting Solutions for Your Industry, Dynamic Ventures Inc., USA). We then divided the number of seeds by the number of spikes to calculate grains per spike.

For secondary cell wall biogenesis (SCWB), samples were collected from the second internode, sliced to expose semi-flat surfaces, and fixed in formaldehyde. Samples were then dehydrated with ethanol and immobilized in paraffin blocks. The stems were cross-sectioned within the blocks using a microtome into 18-20 µm sections. Samples were partially rehydrated and stained with phloroglucinol as described by Pradhan Mitra and Loqué (2014). Samples were then photographed under a Nikon Eclipse NiE microscope using an SD-Ri2 Nikon camera. We then analyzed the samples using an in-house script (see Supplemental Methods S1). The percentage of phloroglucinol-dyed tissue relative to the overall tissue was measured and calculated.

### Genome-Wide Association Analysis

A genome-wide association study (GWAS) was conducted using TASSEL v5.2.5.0 (Bradbury et al., 2007) to identify loci associated with grain yield (GY) under dry and wet environmental conditions in an open field (see field details at (Prusty et al., 2021)). A total of 3,574 SNP markers distributed across the seven barley chromosomes were used for the analysis. Marker–trait associations were evaluated using the General Linear Model (GLM) implemented in TASSEL. To account for population structure, principal component analysis (PCA) was performed, and the resulting principal components were included as covariates in the model. The significance of marker–trait associations was assessed using p-values generated by the GLM. SNPs with −log10(P) ≥ 4.0 were considered significantly associated with the trait. Manhattan plots showing the genomic distribution of marker–trait associations were generated using the online platform SRplot (Tang et al., 2023).

### Time-lapse growth data collection

For time-lapse developmental and physiological analyses, the experiment was conducted in a greenhouse connected to the PlantScreen™ Modular phenotyping platform (PSI, Czech Republic). Here, plants were imaged and analyzed as described by Tietze et al. (2025). Water content was maintained at 55% relative water weight for WW and 25% for WL from the development of the 5^th^ leaf until 65 DAT (days after transplanting) and 20% from 65 DAT until the conclusion of the experiment. Water content was measured daily by pot weight, and watered automatically to match the expected weight.

## Results

### Identification of the diversity among alleles of *Dry2.2*

In previous experiments, we identified that segregation of *Dry2.2* in two HEB-25 families was linked with grain number (GN) maintenance under drought (Merchuk-Ovnat et al., 2018) (Fig. 1A). Considering the HvCEN gene as a strong candidate for the causal diversity underlying these effects (Comadran et al., 2012; Bi et al., 2019; Wang et al., 2020; Göransson et al., 2021), we compared sequence diversity around the gene region to identify potential molecular causality. A G-to-C SNP in exon 4 of HvCEN was previously suggested to underlie drought adaptation (Comadran et al., 2012). However, the wild allele segregating in the HEB-05 family (Barke-Dry2.2^HID065^) lacked this variation, implying that this polymorphism may not be the causal variation we were seeking.

We sequenced several *H. vulgare* (Hv) alleles, including those from cultivars Barke and Noga. These cultivated barley lines served as common backgrounds for the HEB-25 and CMPP populations, respectively (Maurer et al., 2015; Bodenheimer et al., 2025). The wild line used for sequencing was HEB-04-096 (Barke-Dry2.2^HID062^).

Sanger sequencing identified SNPs both upstream and downstream of HvCEN, as well as within the introns of the gene, and a single SNP within the 3’ UTR. We also confirmed the presence of a C-to-G SNP in exon 4, resulting in an Alanine-to-Proline substitution (Fig. 1B). We further identified a 5-base pair indel in the promoter region (679 bp upstream of the translation start site) of the gene. These five base pairs [TTTAT] were lacking in the phenotypically positive plant (Barke-*Dry2.2^Hs_HID062^*) and present in phenotypically negative plants (Barke, Noga). We found that these five base pairs, when present, completed a TATA box. We therefore named this indel CHAT (*Hv**C**EN* **Ha**kuna **T**ata) (Fig. 1C).

### Phenotyping the allelic series of *Dry2.2* as individual plants revealed a specific wild allele that controlled reproductive fitness under drought

Having previously identified that only two out of twenty-five bi-parental populations in the HEB-25 population were associated with drought-resistant grain production (Merchuk-Ovnat et al., 2018), we asked whether these effects are also found among wild alleles originating further south in the Levant, which is characterized by drier barley niches (Chang et al., 2022). We utilized the recently developed wild barley cytonuclear multiparent population (CMPP) (Bodenheimer et al., 2025). Following previous experiments that exhibited maintenance of harvest traits under water-limited (WL) conditions in carriers of the wild *Dry2.2* allele in the Barke background, we performed a similar experiment using the broader genomic panel and its background line, Noga.

The CMPP lines for field experiments were selected based on 50K iSELECT genotyping (Bayer et al., 2017). We confirmed the presence of a wild introgression around *Dry2.2* and winnowed our selection by identifying the direction of the previously reported inversion in this locus (Jayakodi et al., 2020) (Fig. 1A). We further genotyped our selected CMPP lines by HRM to identify the presence or absence of CHAT base pairs (bp) (Fig. 1C) and by TaqMan genotyping for the C-to-G SNP in exon 4 (Table 1). This comparison revealed that the only tested CMPP wild allele carrying CHAT, as found in Barke-*Dry2.2^Hs_HID062^*, originated from B1K-09-07, a wild ecotype from the southern coastal plain (HÜbner et al., 2009).

We selected four CMPP lines (from wild donors B1K-09-07, B1K-29-13, B1K-33-09, and HID386) and backcrossed them into Noga to obtain nearly isogenic advanced backcross fixed siblings at the *Dry2.2* locus (see Methods; Table 1).

Single plants were grown under semi-controlled conditions, classified as well-watered (WW) or water-limited (WL; see Methods). Whole plant phenotype included vegetative dry weight (g), tiller number, spike number, spike weight (g), grain number, and grain dry weight (g). From these, the single spike weight (g), grains per spike, grain dry weight per spike (g), and thousand grain weight (g) were calculated.

Spike weight, single spike weight, grain weight, and grain number were significantly reduced under WL in Barke, Noga, and in Barke-*Dry*2.2^Hv_Barke^ but not in the advanced backcross sibling genotype Barke-*Dry*2.2^Hs_HID062^ (Fig. 2, Tables S2-3). This corroborates our earlier finding that grain number is maintained under WL (Merchuk-Ovnat et al., 2018). None of the other four advanced backcross couplets showed a pattern of trait differentiation between the cultivated and wild carrying lines under WW and WL conditions, except for grain weight between Noga-*Dry*2.2^Hs_HID386^ and Noga-*Dry*2.2^Hv-Noga^ (Fig. S5, Tables S2-3).

**Figure 2.**
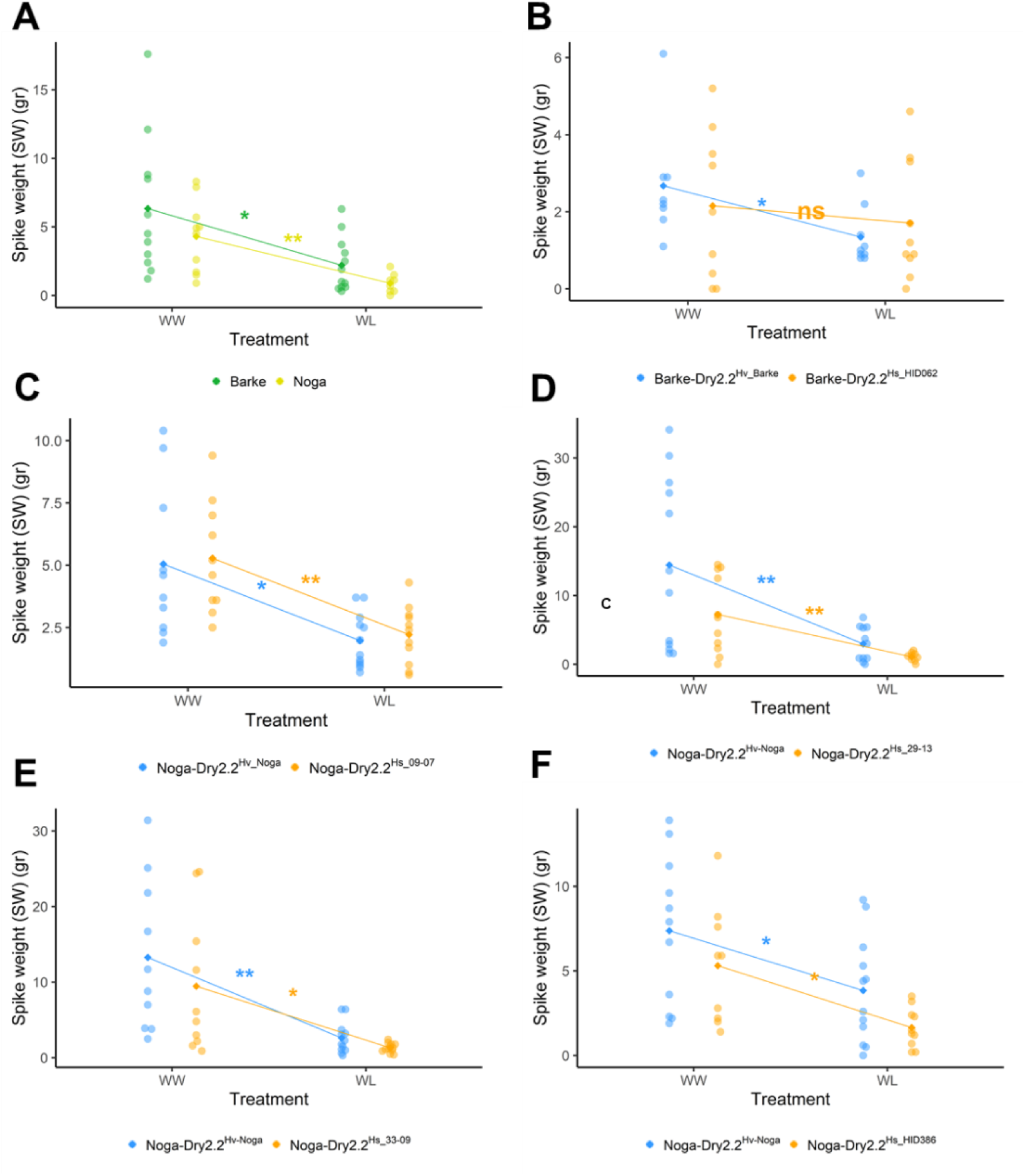
Barke-Dry2.2^HID_062^ alone displayed maintenance of fitness under drought. Spikes were weighed whole in **A)** Controls **B)** HEB-04-096xBarke BC_3_S_1::2_ **C)** B1K-09-07xNoga BC_3_S_1::2_ **D)** B1K-29-13xNoga BC_3_S_1::2_ **E)** B1K-33-09xNoga BC_3_S_1::2_ **F)** HID386 BC_3_S_1::2_. Outliers were removed using the IQR method with a coefficient of 1.5. stars represent significance as calculated by students’ t-test ns=not significant *=<0.05, **<0.01, ***<0.001.

### *Dry2.2* was identified as a cryptic adaptive locus at the plot-level

We wished to examine whether the effects of *Dry*2.2^Hs_HID062^ in the Barke background appeared in the plot, rather than only at the individual level. We held a mini-plot field trial using the HEB-25 population in Gilat, under WW and WL conditions (see Methods). We then performed a GWAS using the same genetic model that previously identified *Dry2.2* in a low-density experiment (Merchuk-Ovnat et al., 2018). We identified several significant marker–trait associations for grain yield under both WL and WW environments (Fig. 3). Under WL conditions, a strong association signal was detected on chromosome 3H. The most significant markers reached −log10(P) values greater than 17, indicating the presence of a major genomic region influencing grain yield under moisture-limited conditions. Additional significant associations were observed on other variables, although with comparatively lower significance levels (Supplementary Table S4).

**Figure 3.**
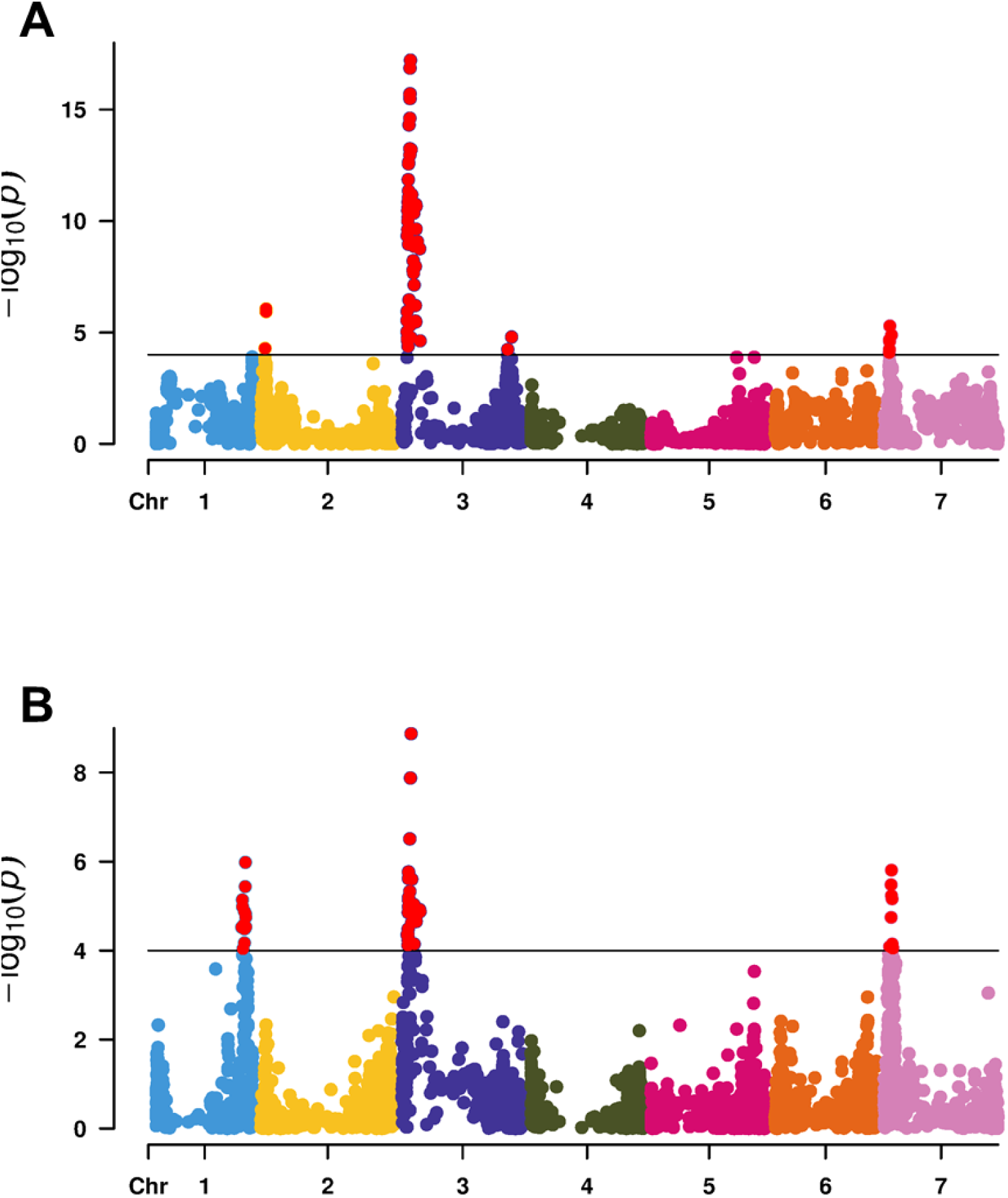
*Dry2.2* remains cryptic in Manhattan plots of GWAS for grain yield under. **A)** dry and **B)** wet conditions. Association analysis was performed using a GLM in TASSEL v5.2.5.0 with 3,574 SNP markers. The x-axis represents chromosome positions, and the y-axis shows −log10(P) values. The horizontal line indicates the significance threshold (−log10(P) = 4.0), and significant SNPs are highlighted in red.

Under WW conditions, significant marker–trait associations were also detected, with the strongest signal again localized to chromosome 3H, where peak markers reached −log10(P) values of approximately 9. Significant loci were additionally identified on chromosomes 1H and 7H, suggesting that these genomic regions contribute to grain yield performance under favorable moisture conditions. Compared with the WL environment, the magnitude and number of significant associations were reduced, indicating a stronger genetic effect of the detected loci under drought stress. Notably, the major locus on chromosome 3H was consistently detected in both environments, suggesting a stable genomic region associated with grain yield across contrasting moisture regimes. Nevertheless, in both WW and WL conditions, neither the *Dry2.2* locus nor any region on chromosome 2 showed a significant effect on GY.

### Differential structural development response to drought linked with the *Dry2.2^Hs_HID062^* wild allele

Identifying the effects of *Dry2.2* in single plants rather than in plots, and focusing primarily on the maintenance of a major fitness trait (grain number), we further asked whether the *Dry2.2* wild alleles had additional effects that could help us understand the mode of action underlying this robustness. Under water-limited conditions, maintaining the structural integrity of the plant’s vasculature through lignification is critical for withstanding high negative water potentials and preventing xylem cavitation (Ntawuguranayo et al., 2024). Thus, we began our exploration of additional phenotypes by focusing on secondary cell-wall biogenesis. Our motivation to investigate this specific trait was further inspired by previous work on the tomato Florigen, the ortholog of the barley HvCEN, which showed that modulation of this gene is intrinsically linked to changes in secondary cell wall biogenesis (SCWB) (Shalit-Kaneh et al., 2019).

Stem samples collected from a segregating H4_96xBarke BC_3_S_1_ population (segregating between Barke-*Dry*2.2^Hv_Barke^ and Barke-*Dry*2.2^Hs_HID062^ types) and grown under WL and WW conditions were stained with phloroglucinol to localize and quantify lignin content (Fig. 4A) (Pradhan Mitra and Loqué, 2014). We also genotyped the same segregating plants by TaqMan for the exon 4 SNP (Table 1). We found that lignin accumulation in Barke-*Dry*2.2^Hv-Barke^ was significantly lower under WL than under WW. In contrast, the lines carrying the wild allele (Barke-*Dry*2.2^Hs-HID062^) showed no significant change in lignin accumulation between WW and WL grown plants (Fig. 4B).

**Figure 4.**
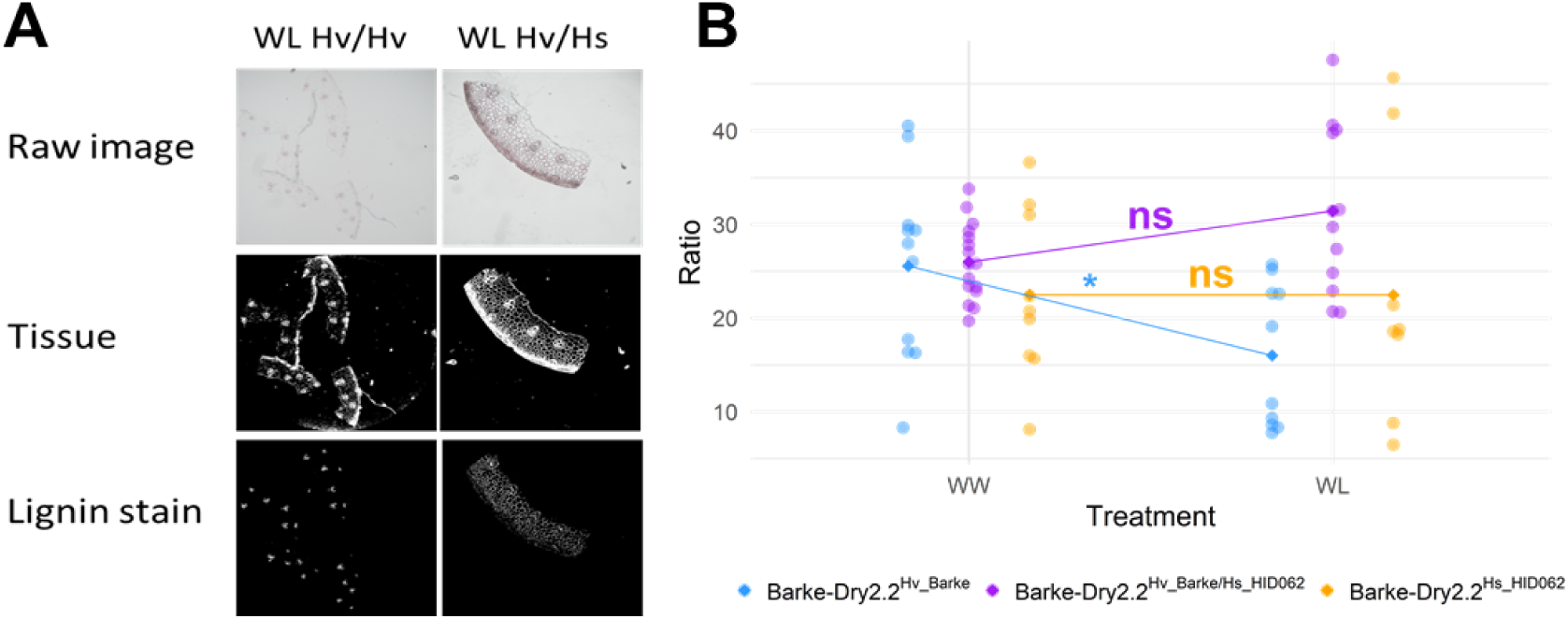
Lignin accumulation in the stem displays a positive correlation with harvest traits. **A)** Samples of images used in analysis. Raw image was taken with a microscope, image of isolated tissue generated by an in-house script, and image of isolated lignin stain generated by an in-house script (supplementary Methods S1). **B)** Stained tissue ratio was calculated by the number of white pixels in the lignin stain image divided by the number of white pixels in the tissue image. Outliers identified using the SD method, with a threshold of 2. Stars indicate significance as calculated by students’ t-test.

### Time-lapse phenotyping revealed pleiotropic effects of *Dry2.2*^Hs_HID062^ on growth

To gain a further understanding of the function of *Dry2.2*, we needed to move beyond end-of-life yield metrics (“final phenotype”) and capture the transient, dynamic physiological shifts that single-point harvest misses. We hypothesized that the wild and cultivated alleles dictate divergent physiological strategies, such as stomatal regulation, canopy cooling, and the initiation of senescence, immediately upon the onset of water limitation.

To non-destructively capture these early temporal interactions at a high resolution, we performed an automated time-lapse using the PlantScreen™ Modular phenotyping platform (PSI, Czech Republic) (Abdelhakim et al., 2024; Tietze et al., 2025). Single plants were grown in pots under semi-controlled conditions and were considered well-watered (WW) or water-limited (WL). Images were taken at various time points over 94 days. To assess the impact of *Dry2.2* on plant responses to WL over time, we used multiple imaging sensors, including RGB, thermal infrared (IR), chlorophyll fluorescence, and hyperspectral imaging (VNIR, SWIR). We focused on RGB and hyperspectral imaging (SWIR), as the former emerged as an important predictor of plant yield success (Tietze et al., 2025), and the latter directly reflects plant water status.

### Wild allele introgressions maintained higher water content under WL

As drought leads to a water deficit, we examined leaf water content. Water content index was calculated as a reflectance ratio of 1440 nm to 960 nm. 1440 nm is strongly absorbed by water. When reflectance decreases at 1440 nm, resulting in a lower water index value, it indicates a well-hydrated leaf. In contrast, when reflectance increases at 1440 nm, resulting in a higher index value, it indicates a more desiccated leaf.

Water Content Index (WCI) was ascertained using hyperspectral imaging (SWIR). Barke-*Dry*2.2^Hs_HID062^, Barke-*Dry*2.2^Hv_Barke^, and Barke displayed a significantly elevated WCI in WL when compared to WW, specifically between 34-55 DAT (Tukey-Kramer HSD test p-value< 2.22E-308, <2.22E-308, and 1.9E-13, respectively) (Fig. 5). Under WW, Barke-*Dry*2.2^Hs_HID062^ had elevated WCI when compared to Barke-*Dry*2.2^Hv_Barke^ and Barke (p-value=3.77e-9 and 6.44e-15, respectively). However, WW Barke-*Dry*2.2^Hs_HID062^ was comparable to the cultivated WL lines. In WL, WCI was significantly elevated in Barke-*Dry*2.2^Hs_HID062^ compared to Barke-*Dry*2.2^Hv_Barke^ and Barke (p-value=1.32E-10 and < 2.22E-308, respectively) (Fig. 5B).

**Figure 5.**
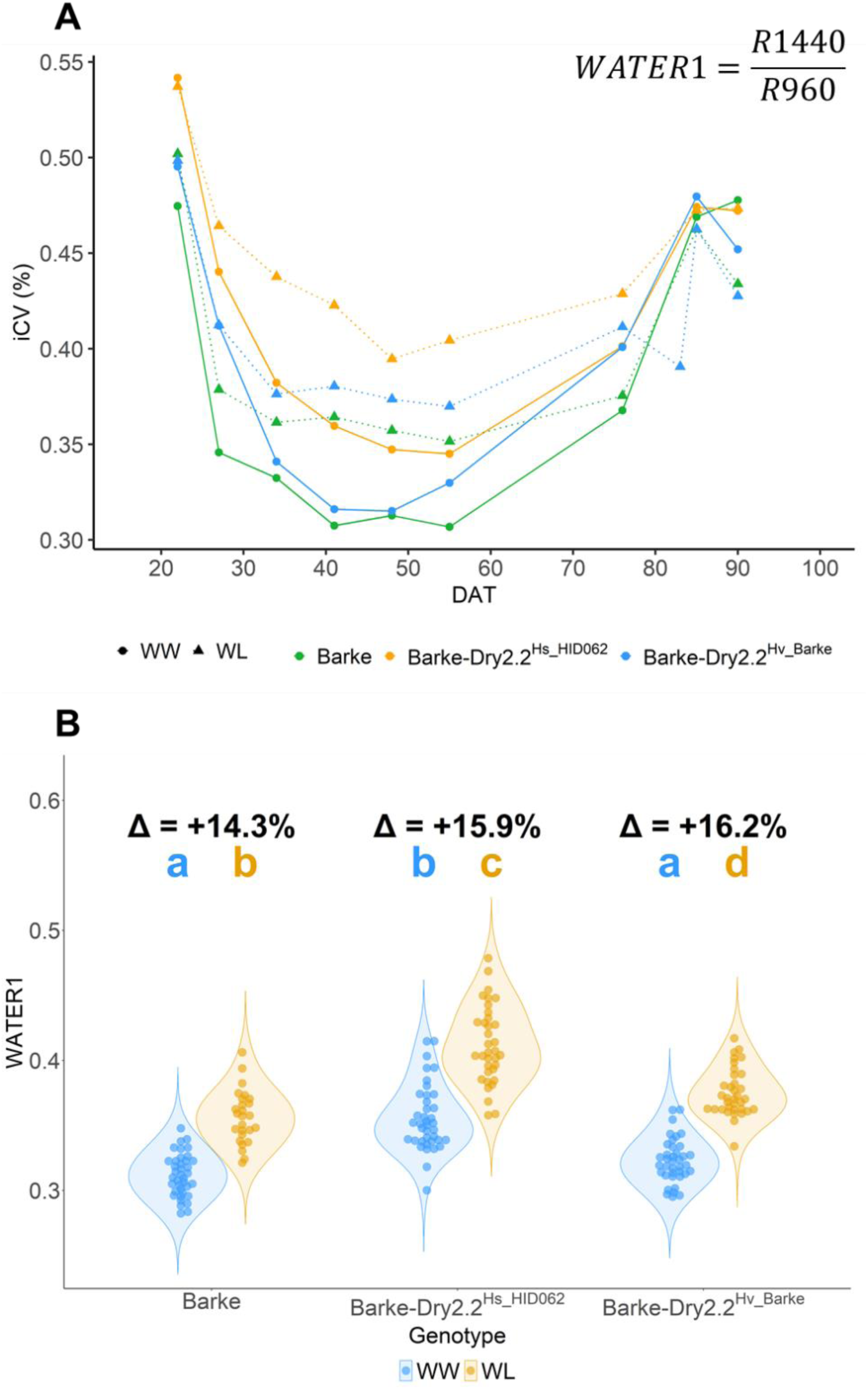
Water content index (WCI) was elevated in the wild allele and WL. **A)** Average of WCI across whole experiment (DAT 20-90) **B)** Results DAT 34-55. Δ signifies the percentage of difference between the averages of the different treatments for each line. Outliers were removed using the IQR method with a coefficient of 1.5. Letters signify significant differences as calculated by the Tukey-Kramer HSD test.

Taken together, these results implied a lower water content in Barke-*Dry*2.2^Hs_HID062^, even under WW conditions, compared with its cultivated counterparts. Furthermore, the water content of the wild allele line in WW conditions was comparable to that found in the cultivated allele lines in WL.

### The senescence rate was maintained in HsDry2.2^HID062^ introgression line under drought

Having previously identified that the wild allele is linked with delayed senescence under semi-controlled experiments (Merchuk-Ovnat et al., 2018), we wished to re-examine these effects in the PSI platform at a higher time resolution. Senescence, as the loss of green pigmentation in the leaves of plants, is a natural end of barley’s life cycle. The onset and rate of senescence are expected to be affected by drought (Chibane et al., 2021). Based RGB imaging, we analyzed the percentage of green plant tissue (RGB (68,85,63)) relative to total tissue and found, as expected, that it declined nearly linearly after ∼50 DAT (Fig. 6A). This rate was consistent across the lines grown in WW conditions (Barke, Barke-*Dry*2.2^Hv_Barke,^ and Barke-*Dry*2.2^Hs_HID062^ rates −1.58, −1.51, and −1.61, respectively). However, in WL, both the Barke and Barke-*Dry*2.2^Hv_Barke^ displayed a significantly milder slope (slope=-1.22, ANCOVA DAT x Treatment p-value=3.01E-016 and slope=-1.25, p-value=5.47E-09, respectively), which did not occur in Barke-*Dry*2.2^Hs_HID062^ (slope=-1.57, t-test p-value=0.79) (Fig. 6B).

**Figure 6.**
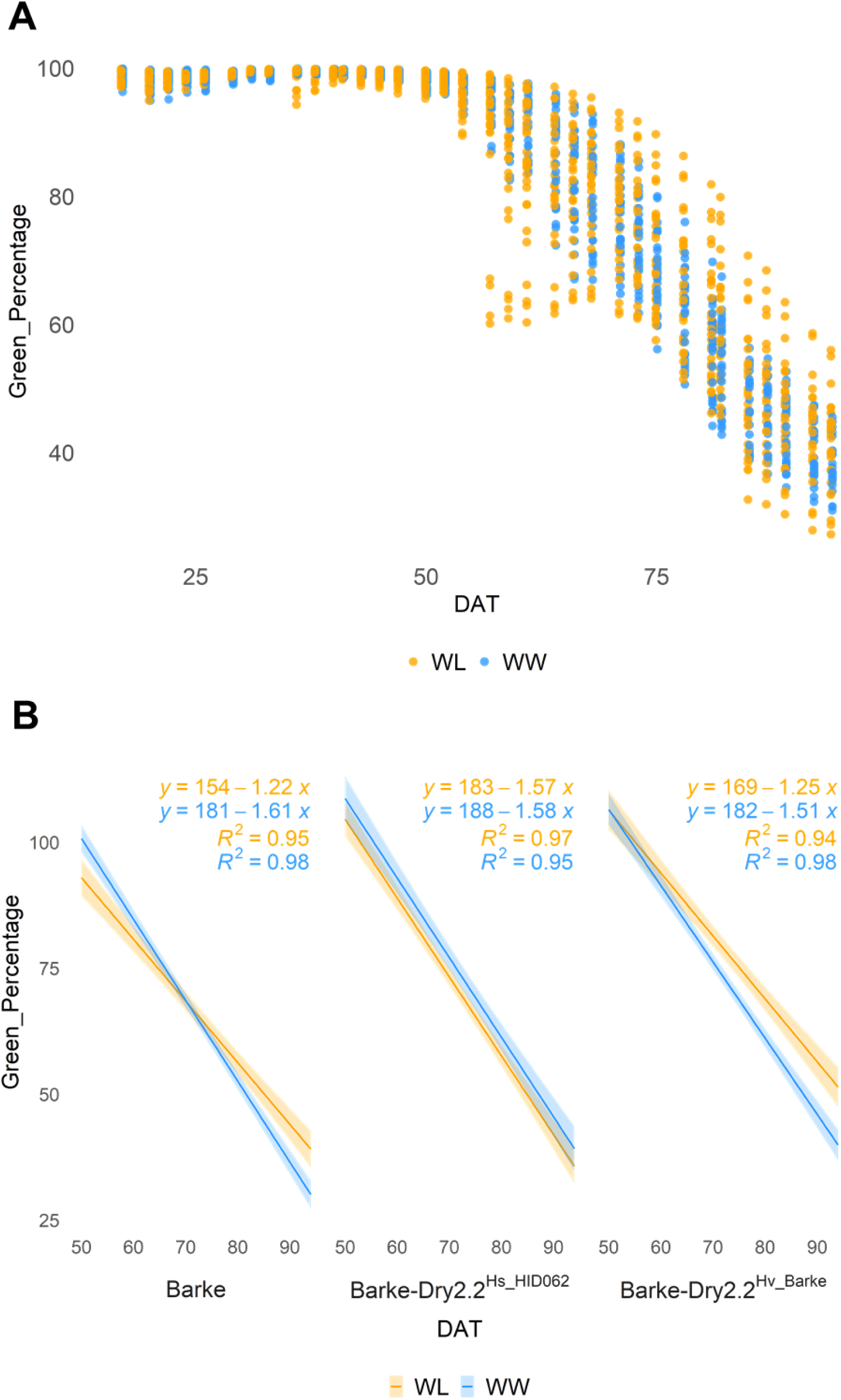
Senescence rate was maintained in wild allele under WL. **A)** percentage of plant in shades of green (RGB (68,85,63)) **B)** linear regression of senescence starting at 50 DAT. Outliers were removed using the IQR method with a coefficient of 1.5. Equations were calculated from averages, with the R^2^ indicating the goodness of the linear fit.

### Dry2.2^Hs_HID062^ advances flowering under drought

*Dry2.2* is involved in the timing of life events, such as flowering. We wanted to see whether, under our experimental conditions, which took place in another country and with longer days than in the previous study (see Methods) (Merchuk-Ovnat et al., 2018), the results would differ. Thus, we identified the heading time as the DAT at which the first awns emerged.

While both Barke-*Dry*2.2^Hv_Barke^ and Barke-*Dry*2.2^Hs_HID062^ displayed a significant advance in heading time compared to Barke in both WW (p-value=6.54E-7 and 7.08E-8, respectively) and WL conditions (p-value=1.24E-7 and < 2.22E-308, respectively), only the latter exhibited an advance under WL, when compared to WW (p-value=1.27e-5) (Fig. 7A). This contrasted with earlier findings showing no GxE, where flowering time in *Dry2.2* lines was reported to be identical under both environments (Merchuk-Ovnat et al., 2018).

**Figure 7.**
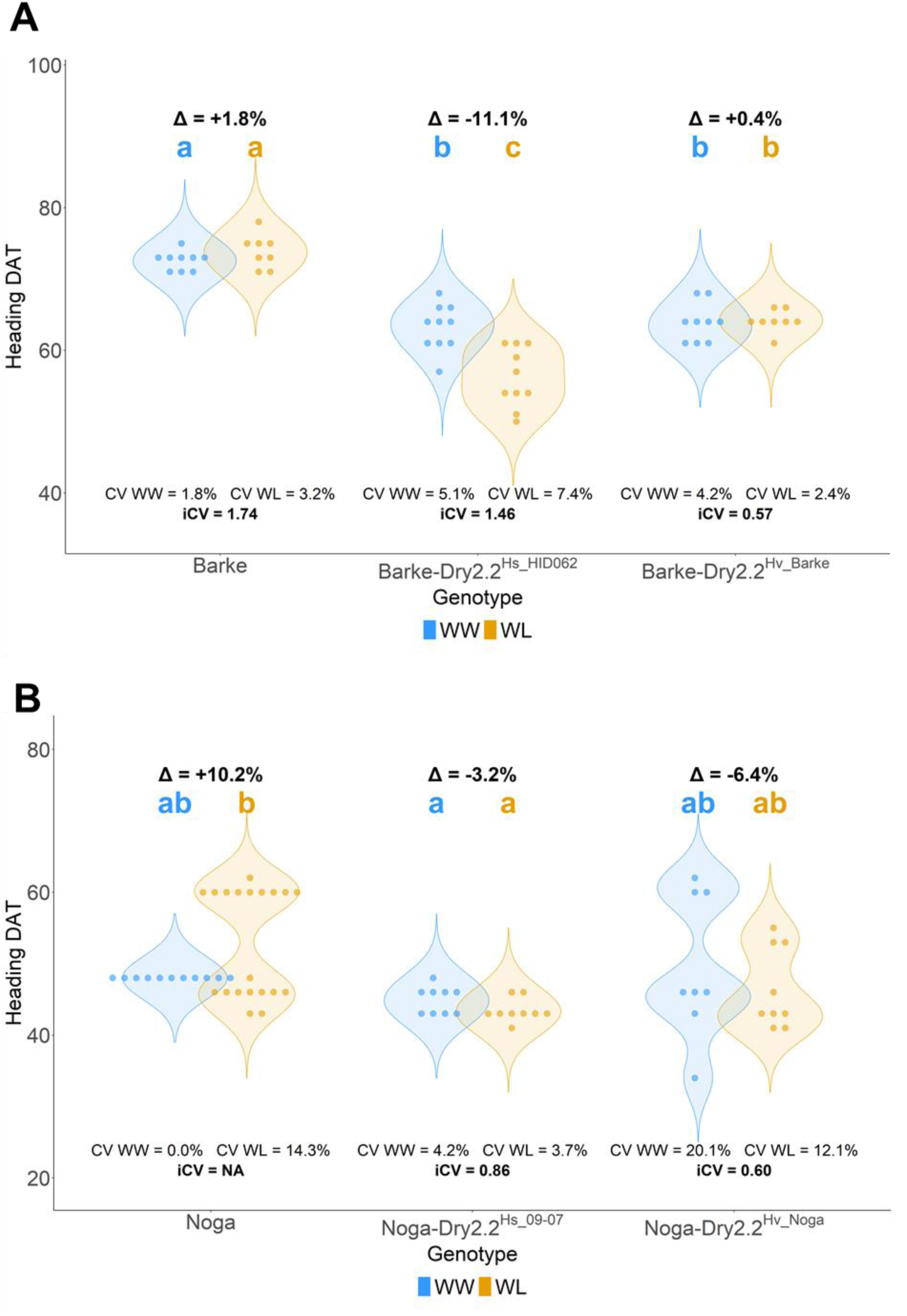
Plant Heading Date is affected by WL. **A)** Heading DAT in Barke and HEB-04-096 derived lines **B)** Heading DAT in Noga and B1K-09-07 derived lines. Δ signifies the percentage of difference between the averages of the different treatments for each line. Outliers were removed using the IQR method with a coefficient of 1.5. Letters signify significant differences as calculated by the Tukey-Kramer HSD test.

As seen previously in the harvest trait analysis (Fig. 2B), heading time in Noga-Dry2.2^Hs_09-07^ behaved like the cultivated lines, rather than Barke-*Dry*2.2^Hs_HID062^. No significant advances in heading DAT were noted in any line, under any condition (Fig. 7B).

While examining the distribution of heading time, we observed that variance differed across genetic backgrounds. In the Noga background, we observed a significant difference between carriers of the wild (Noga-Dry2.2^Hs_09-07^) and cultivated (Noga-Dry2.2^Hv_Noga^) alleles. Under both environments, carriers of the cultivated allele showed a clear expansion in trait distribution, exhibiting a much higher coefficient of variation (CV) for heading DAT when compared to carriers of the wild allele (20.1% vs. 4.2% for WW and 12.1% vs. 3.7% for WL) (Fig. 7B).

Interestingly, this profound reduction in phenotypic variance conferred by the wild allele did not manifest in the Barke background; the CV of Barke-Dry2.2^Hs_HID062^ did not show a lower CV compared to Barke-*Dry*2.2^Hv_Barke^ for Heading DAT (Fig. 7A).

### Dry2.2^HID062^ was negatively correlated with inter-plant variation

The observed differences in the distribution of individuals’ maturity values prompted us to more closely examine the growth dynamics of the different plant populations. An obvious expectation is that plant volume can also be significantly influenced by drought, as with less water, plants are less likely to invest in biomass production. We therefore calculated the plant volume by analyzing RGB images from side and top views. This analysis was performed on the advanced backcross pairs with wild alleles of HID062 or B1K-09-07, compared with their genetically related control lines.

As expected, plant volume was larger under WW conditions than under WL (Fig. 8A, B). Nevertheless, an exploration of volume variation revealed a higher coefficient of variation (CV) in cultivated allele-carrying lines in WL, which more than doubled that of wild allele-carrying lines. In WL, by 27 DAT, the cultivated line Barke-*Dry*2.2^Hv_Barke^ had a 200% higher CV than Barke-Dry2.2^Hs_HID062^ (36.1% vs 17.5%, respectively) (Fig 8C). Similarly, in WL, the Noga-Dry2.2^Hv_Noga^ genotype had twice the CV of Noga-Dry2.2^Hs_09-07^ at 40 DAT (38.3% vs. 19.3%, respectively) (Fig. 8D).

**Figure 8.**
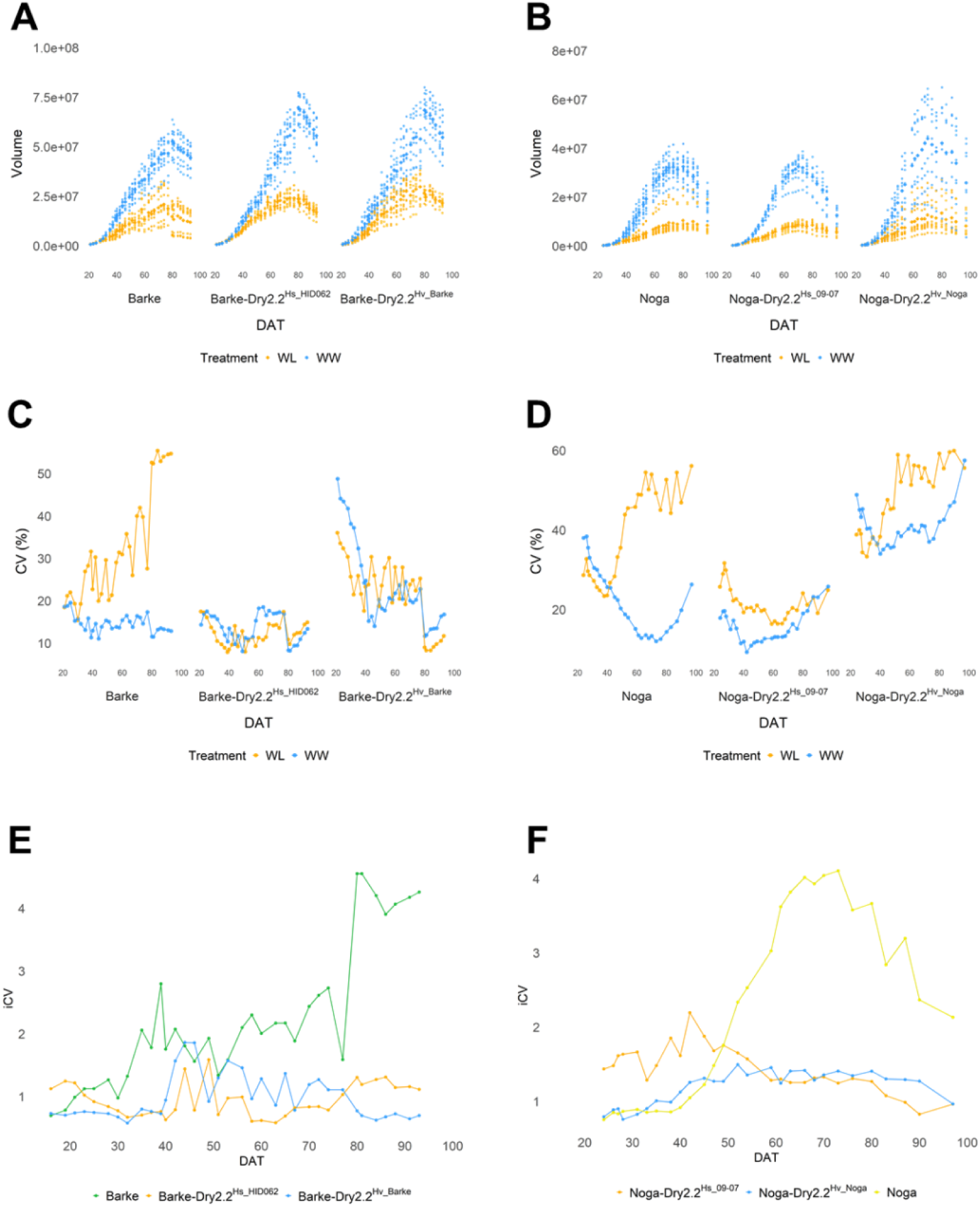
Plant Volume shows wider distribution between individuals in cultivated *Dry2.2* carrying lines under WL. **A)** Volume in Barke and HEB-04-096 derived lines **B)** Volume in Noga and B1K-09-07 derived lines **C)** Volume iCV in Barke and HEB-04-096 derived lines **D)** Volume CV in Noga and B1K-09-07 derived lines **E)** Volume iCV (WL CV/WW CV) in Barke and HEB-04-096 derived lines **F)** Volume iCV (WL CV/WW CV) in Noga and B1K-09-07 derived lines.

Since both segregating advanced backcross lines, either carrying the wild or cultivated allele of *Dry2.2*, could carry additional segregating loci between individuals, we also explored the variation in the fully cultivated inbred lines. Under WL, by 35 DAT Barke had more than doubled the CV of *Dry*2.2^Hs_HID062^ (26.9% and 9.72%) (Fig. 8C) and at 52 DAT Noga had doubled the CV of the wild allele carrier (43.91% and 19.63%) (Fig. 8D). Profoundly, both plant populations of Noga and Barke displayed a much higher iCV (WL CV/WW CV) (Fridman, 2015) than their wild introgression counterparts, starting at DAT ∼50 (Fig. 8E, F).

## Discussion

### Identification of Possible Causal Variations at the *Dry2.2* locus and GxG Interactions

In our search for the molecular causality underlying the maintenance of fitness under water deficit, we focused on *HvCEN* as a strong candidate gene due to its orthology with plant architecture and flowering genes (Comadran et al., 2012a; Shalit-Kaneh et al., 2018; Bi et al., 2019). Sequencing revealed several polymorphisms, including a non-synonymous SNP in exon 4 and a specific indel in the promoter region (CHAT) (Fig. 1). While these polymorphisms helped us locate similar lines, such as *Dry2.2^Hs_09-07^*, this specific introgression did not maintain grain production under drought conditions, though it did correlate with differential phenotypic uniformity in other traits (Fig. 7B, 8B, D, F).

This partial phenotypic replication suggests that *HvCEN* polymorphisms do not act alone; rather, the locus’s causal output is heavily dependent on a dynamic interplay of genomic background and environmental triggers. The observation that the reduction in phenotypic variance for maturity traits is prominent in the Noga background but absent in the Barke background (Fig. 7) implies that the full effect of the *Dry2.2* locus requires specific genotype-by-genotype (GxG) epistatic interactions with the broader cultivated genome.

Furthermore, our detailed comparisons during the European fall-to-winter period reveal that *Dry2.2* penetrance relies on genotype-by-environment (GxE) interactions. For example, Barke-Dry2.2^Hs_HID062^ displayed a significant advance in heading time under water-limited conditions compared to well-watered conditions (Fig. 7), an active stress-escape response that contrasts with previous studies showing identical heading effects across environments (no GxE) under Mediterranean winter (Merchuk-Ovnat et al., 2018). Similarly, the previously reported effects on senescence in a pot experiment in the Mediterranean (Merchuk-Ovnat et al., 2018) differed from the European experiments at PSI (Fig. 6).

Therefore, *Dry2.2* functions not as a static “on/off” switch, but as a complex developmental dial whose output is dictated by local polymorphisms, the recipient genome, and active environmental triggers.

### Mechanical and Canopy Rigor: The Structural Basis of Canalization

The structural robustness supporting the wild allele’s canalized escape strategy is grounded in the maintenance of secondary cell wall biogenesis (SCWB) (Fig. 4). Under water-limited conditions, maintaining the structural integrity of cereals’ vasculature, partially through lignification, is critical for withstanding high negative water potentials and preventing xylem cavitation (Ntawuguranayo et al., 2024). Our histochemical evaluations revealed that stable accumulation of stem lignin in *Dry2.2^Hs-HID062^* carriers protects the vasculature under stress (Fig. 4), allowing the wild introgression lines to maintain seed production. Furthermore, we observed a preemptive response in *Dry2.2^Hs-HID062^* carriers regarding water content in the canopy (Fig. 5). The water content in *Dry2.2^Hs-HID062^* carriers in WW is significantly lower than that in *Dry2.2^Hv-Barke^* carriers in WW, and comparable to WL *Dry2.2^Hv-Barke^*carriers. This lower water content suggests that the plant is primed to function under low water potentials, such as those found during drought. Conversely, the cultivated lines (Barke and Noga) exhibit developmental non-canalization, characterized by a significant reduction in stem lignin accumulation under stress.

### Domestication and the Paradoxical Selection for Bet-Hedging under Stress

Agricultural domestication is traditionally viewed as a systematic sweep toward phenotypic uniformity, with heavy selection for canalized phenotypes. Our discovery that the cultivated allele (*Dry2.2^Hv^*) actively fosters a “bet-hedging” risk-spreading strategy under water limitation challenges this linear paradigm (Fig 7-8). In highly volatile environments where the onset, duration, and severity of drought are unpredictable, a uniform developmental response can be evolutionarily catastrophic for a dense population. By generating a diverse array of individual developmental trajectories (expanding the coefficient of variation for traits like volume and maturity) from a single genotype, the cultivated population ensures that at least a subset of individuals will successfully set seed regardless of when environmental stress peaks.

This heterogeneity explains a critical mapping paradox observed in standard plot-level agricultural evaluations. Standard genome-wide association studies (GWAS) are inherently mean-centric; they mathematically rely on discovering directional shifts in a plot’s central tendency. Because the cultivated allele’s primary adaptive effect under stress is to expand inter-individual variance rather than shifting the plot-level average, its distinct drought-responsive signal is statistically masked. In contrast, these same central-tendency models consistently capture loci like *DOC3* on Chromosome 3H (Prusty et al., 2021), which optimize macro-environmental grain yield through distinct, mean-shifting adaptive sweeps (Fig. 3).

However, the adaptive value of this variance-driven strategy becomes strikingly apparent when shifting the lens from short-term uniform agricultural plots to long-term evolutionary competition. This provides a critical missing link to the selection signatures identified in the California Composite Cross (CC) (Landis et al., 2024). In this century-scale experiment, barley populations were grown in high-density, highly competitive large plots under a volatile Mediterranean climate. While selection on *HvCEN* in the CC was previously attributed primarily to shifts in average flowering time, our findings suggest the true evolutionary advantage of the cultivated allele in such environments is its facilitation of group-level bet-hedging. In a dense, competing stand facing unpredictable drought (typical to CA or Mediterranean climate), intentionally spreading developmental risk ensures that a subset of individuals will successfully set higher amounts of seed regardless of when the stress peaks. Thus, while standard plot evaluations successfully map loci like *DOC3* for short-term (Prusty et al., 2021), uniform yield maximization, long-term competitive environments actively select for the cultivated allele of *Dry2.2* to ensure population survival through resilient, risk-spreading heterogeneity.

### Epistasis, Structural Inversions, and the Future of Resilient Crop Selection

The contrasting penetrance of these phenotypic variances across different cultivated backgrounds confirms that *Dry2.2* is deeply contingent upon the broader recipient genome. This epistatic masking, coupled with a local structural inversion that completely suppresses local recombination (Fig. 1) (Comadran et al., 2012), has effectively locked the wild and cultivated alleles into tightly bound haplotype blocks. Consequently, determining whether the observed suite of resilience traits is driven by true pleiotropy of a single master regulator (*HvCEN*) or by a cluster of tightly linked genes remains an outstanding challenge.

To resolve this genetic entanglement, ongoing experimental efforts are focused on breaking this linkage suppression by transferring the *Dry2.2* introgression into cv. Morex (Rasmusson and Wilcoxson, 1979). Because cv. Morex lacks the structural inversion at this specific locus (Jayakodi et al., 2020) it can provide a definitive genetic test to uncouple *HvCEN* from its neighboring passenger loci.

Ultimately, as agricultural systems face increasingly volatile climates, crop improvement must move beyond the narrow constraint of absolute uniformity. Unlocking the genetic dials that control developmental canalization versus bet-hedging will allow breeders to actively design future crop populations by choosing between uniform individual robustness or resilient, risk-spreading population heterogeneity, depending on target environmental predictability.

## Acknowledgements

This study in the framework of the CAPILTALISE project was funded by the European Union’s Horizon 2020 research and innovation program (grant no. 862201). We thank Hani Zemach (Microscopy and Imaging, Volcani Institute) for her assistance in sectioning and staining samples for lignin quantifications.

## Competing interests

The authors declare no competing interests

## Author contributions

A.K.S. designed and developed the crosses, organized the phenotypic and genotypic datasets for the greenhouse and PSI experiments, wrote the script for image analysis if lignification, analyzed the data, and wrote the initial draft and the final version of the manuscript with E.F.; M.R.P. organized the phenotypic and marker dataset for the mini-plot experiments, analyzed the data, and wrote the initial draft for the GWAS; L.A. and K.P. managed and obtained raw data of experiments on the PlantScreen phenomics platform. A.B. managed the seeds for the field trials and assisted with phenotyping; E.F. supervised the data analysis, wrote the initial draft and final version of the paper with A.K.S. All co-authors critically reviewed the manuscript

## Data availability

All raw and pruned datasets are available in the supplementary material. The custom Python script used for image analysis of lignification is fully accessible at [https://github.com/AyeletKS/Lignification-calculation/blob/main/image.py]

## Supporting Information

**Note:** Table S1, Table S2, and Methods S1 are included below in this document. Data Tables S3 through S11 are hosted externally in the file **AKS_Dry2.2 _SupTable_170626.xlsx**

The following Supporting Information is available for this article:

**Table S1.**
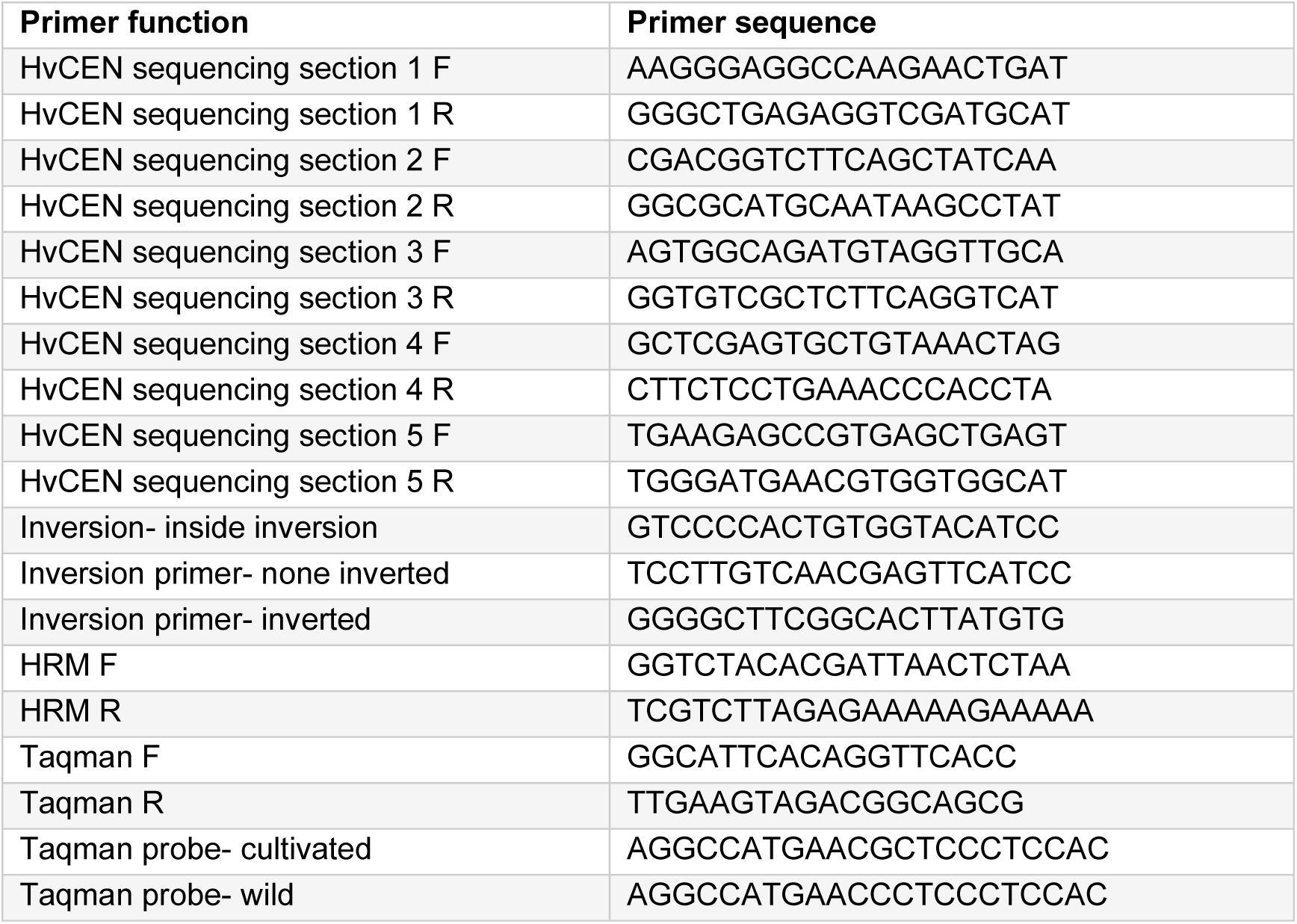
Primers used in study.

**Table S2.** Changes in Harvest Traits Between Control and Drought Conditions. Outliers identified with interquartile range (IQR) method, with a threshold of 1.5 Stars and colors indicate significance.

|  | Overall weight | Number of tillers (TN) | Number of spikes (SN) | Spike weight (SW) | Single spike weight (SSW) | Grain weight (GW) | Single spike grain weight (SSGW) | Grain number (GN) | Grains per spike (GPS) | 1000 grain weight |
| --- | --- | --- | --- | --- | --- | --- | --- | --- | --- | --- |
| **Barke** | * | * | ** | * | * | * | NS | ** | NS | NS |
| **Barke-Dry2.2<sup>Hv</sup>-Barke** | NS | NS | NS | ** | ** | *** | *** | * | * | NS |
| **Barke-Dry2.2<sup>Hs</sup>-HID062** | NS | NS | NS | NS | NS | NS | NS | NS | NS | NS |
| **Noga** | ** | NS | NS | ** | ** | * | NS | ** | * | ** |
| **Noga-Dry2.2<sup>Hv</sup>-Noga** | * | *** | * | * | NS | * | * | * | * | NS |
| **Noga-Dry2.2<sup>Hs</sup>-09-07** | ** | NS | *** | ** | NS | ** | NS | ** | NS | NS |
| **Noga-Dry2.2<sup>Hv</sup>-Noga** | ** | ** | ** | ** | NS | * | NS | * | NS | NS |
| **Noga-Dry2.2<sup>Hs</sup>-29-13** | ** | ** | ** | ** | *** | ** | *** | ** | ** | * |
| **Noga-Dry2.2<sup>Hv</sup>-Noga** | ** | ** | ** | ** | ** | ** | * | ** | NS | NS |
| **B1K-Noga-Dry2.2<sup>Hs_33-09</sup>** | * | ** | ** | * | NS | ** | NS | ** | NS | NS |
| **Noga-Dry2.2<sup>Hv-Noga</sup>** | * | ** | NS | * | NS | ** | * | ** | NS | * |
| **Dry2.2<sup>Hs_HID386</sup>** | * | * | * | * | NS | NS | * | * | * | NS |

**Table S3** Field trial harvest traits

**Table S4** GWAS results

**Table S5** Lignin ratios

**Table S6** WCI results PSI Barke-HEB_04_096 BG

**Table S7** Green percentage PSI Barke-HEB_04_096 BG

**Table S8** Heading DAT PSI Barke-HEB_04_096 BG

**Table S9** Heading DAT PSI Noga-B1K_09_07 BG

**Table S10** Volume results PSI Barke-HEB_04_096 BG

**Table S11** Volume results PSI Noga-B1K_09_07 BG

**Methods S1** Code for relative lignification [https://github.com/AyeletKS/Lignification-calculation/blob/main/image.py]

## Notes

https://github.com/AyeletKS/Lignification-calculation/blob/main/image.py

### Competing Interest Statement

The authors have declared no competing interest.

## Bibliography

Abdelhakim, L. O. A., Pleskačová, B., Rodriguez-Granados, N. Y., Sasidharan, R., Perez-Borroto, L. S., Sonnewald, S., Gruden, K., Vothknecht, U. C., Teige, M., and Panzarová, K. (2024). High throughput image-based phenotyping for determining morphological and physiological responses to single and combined stresses in potato. J Vis Exp Advance Access published June 7, 2024, doi:10.3791/66255.

Abley, K., Goswami, R., and Locke, J. C. W. (2024). Bet-hedging and variability in plant development: seed germination and beyond. Philos Trans R Soc Lond B Biol Sci 379:20230048.

Alpert, P., and Simms, E. L. (2002). The relative advantages of plasticity and fixity in different environments: When is it good for a plant to adjust? Evolutionary Ecology 16:285–297.

Alseekh, S., Tong, H., Scossa, F., Brotman, Y., Vigroux, F., Tohge, T., Ofner, I., Zamir, D., Nikoloski, Z., and Fernie, A. R. (2017). Canalization of tomato fruit metabolism. The Plant Cell 29:2753–2765.

Badr, A., Müller, K., Schäfer-Pregl, R., El Rabey, H., Effgen, S., Ibrahim, H. H., Pozzi, C., Rohde, W., and Salamini, F. (2000). On the origin and domestication history of Barley (*Hordeum vulgare*). Mol Biol Evol 17:499–510.

Bayer, M. M., Rapazote-Flores, P., Ganal, M., Hedley, P. E., Macaulay, M., Plieske, J., Ramsay, L., Russell, J., Shaw, P. D., Thomas, W., et al. (2017). Development and evaluation of a barley 50k iSelect SNP array. Front Plant Sci 8:1792.

Beche, E., Gillman, J. D., Song, Q., Nelson, R., Beissinger, T., Decker, J., Shannon, G., and Scaboo, A. M. (2020). Nested association mapping of important agronomic traits in three interspecific soybean populations. Theor Appl Genet 133:1039–1054.

Bi, X., Esse, W. V., Mulki, M. A., Kirschner, G., Zhong, J., Simon, R., and Korff, M. V. (2019). CENTRORADIALIS interacts with FLOWERING LOCUS T-like genes to control floret development and grain number. Plant Physiology 180:1013–1030.

Bodenheimer, S., Bdolach, E., Be’ery, A., Tiwari, L. D., Perez-Alfaro, R. S., Yang, S., Koenig, D., and Fridman, E. (2025). Harnessing cytonuclear diversity to map barley spike traits using the cytonuclear multi-parent population. Genetics 231:iyaf167.

Bradbury, P. J., Zhang, Z., Kroon, D. E., Casstevens, T. M., Ramdoss, Y., and Buckler, E. S. (2007). TASSEL: software for association mapping of complex traits in diverse samples. Bioinformatics 23:2633–2635.

Bradley, D., Carpenter, R., Copsey, L., Vincent, C., Rothstein, S., and Coen, E. (1996). Control of inflorescence architecture in Antirrhinum. Nature 379:791–797.

Brooker, R., Brown, L. K., George, T. S., Pakeman, R. J., Palmer, S., Ramsay, L., Schöb, C., Schurch, N., and Wilkinson, M. J. (2022). Active and adaptive plasticity in a changing climate. Trends in Plant Science 27:717–728.

Chang, C.-W., Fridman, E., Mascher, M., Himmelbach, A., and Schmid, K. (2022). Physical geography, isolation by distance and environmental variables shape genomic variation of wild barley (*Hordeum vulgare* L. ssp. *spontaneum*) in the Southern Levant. Heredity (Edinb) 128:107–119.

Chibane, N., Caicedo, M., Martinez, S., Marcet, P., Revilla, P., and Ordás, B. (2021). Relationship between delayed leaf senescence (stay-green) and agronomic and physiological characters in maize (Zea mays L.). Agronomy 11:276.

Comadran, J., Kilian, B., Russell, J., Ramsay, L., Stein, N., Ganal, M., Shaw, P., Bayer, M., Thomas, W., Marshall, D., et al. (2012). Natural variation in a homolog of Antirrhinum CENTRORADIALIS contributed to spring growth habit and environmental adaptation in cultivated barley. Nature genetics 44:1388–92.

de Groot, D. H., Tjalma, A. J., Bruggeman, F. J., and van Nimwegen, E. Effective bet-hedging through growth rate dependent stability. Proc Natl Acad Sci U S A 120:e2211091120.

Debat, V., and David, P. (2001). Mapping phenotypes: canalization, plasticity and developmental stability. Trends in Ecology & Evolution 16:555–561.

Fridman, E. (2015). Consequences of hybridization and heterozygosity on plant vigor and phenotypic stability. Plant Science 232:35–40.

Göransson, M., Sigurdardottir, T. H., Lillemo, M., Bengtsson, T., and Hallsson, J. H. (2021). The winter-type allele of HvCEN Is associated with earliness without severe yield penalty in Icelandic spring barley (*Hordeum vulgare* L.). Frontiers in Plant Science 12:1801.

Hanano, S., and Goto, K. (2011). Arabidopsis TERMINAL FLOWER1 is involved in the regulation of flowering time and inflorescence development through transcriptional repression. Plant Cell 23:3172–3184.

HÜbner, S., HÖffken, M., Oren, E., Haseneyer, G., Stein, N., Graner, A., Schmid, K., and Fridman, E. (2009). Strong correlation of wild barley (*Hordeum spontaneum*) population structure with temperature and precipitation variation. Molecular Ecology 18:1523–1536.

Jayakodi, M., Padmarasu, S., Haberer, G., Bonthala, V. S., Gundlach, H., Monat, C., Lux, T., Kamal, N., Lang, D., Himmelbach, A., et al. (2020). The barley pan-genome reveals the hidden legacy of mutation breeding. Nature Advance Access published 2020, doi:10.1038/s41586-020-2947-8.

Lachowiec, J., Queitsch, C., and Kliebenstein, D. J. (2016). Molecular mechanisms governing differential robustness of development and environmental responses in plants. Ann Bot 117:795–809.

Laitinen, R. A. E., and Nikoloski, Z. (2019). Genetic basis of plasticity in plants. J Exp Bot 70:739–745.

Landis, J. B., Guercio, A. M., Brown, K. E., Fiscus, C. J., Morrell, P. L., and Koenig, D. (2024). Natural selection drives emergent genetic homogeneity in a century-scale experiment with barley. Science 385:eadl0038.

Lempe, J., Lachowiec, J., Sullivan, A. M., and Queitsch, C. (2013). Molecular mechanisms of robustness in plants. Curr Opin Plant Biol 16:62–69.

Maurer, A., Draba, V., Jiang, Y., Schnaithmann, F., Sharma, R., Schumann, E., Kilian, B., Reif, J. C., and Pillen, K. (2015). Modelling the genetic architecture of flowering time control in barley through nested association mapping. BMC Genomics 16.

Merchuk-Ovnat, L., Silberman, R., Laiba, E., Maurer, A., Pillen, K., Faigenboim, A., and Fridman, E. (2018). Genome scan identifies flowering-independent effects of barley HsDry2.2 locus on yield traits under water deficit. Journal of Experimental Botany 69.

Mitchell, J., Johnston, I. G., and Bassel, G. W. (2017). Variability in seeds: biological, ecological, and agricultural implications. J Exp Bot 68:809–817.

Ntawuguranayo, S., Zilberberg, M., Nashef, K., Bonfil, D. J., Bainsla, N. K., Piñera-Chavez, F. J., Reynolds, M. P., Peleg, Z., and Ben-David, R. (2024). Stem traits promote wheat climate-resilience. Front Plant Sci 15:1388881.

Pradhan Mitra, P., and Loqué, D. (2014). Histochemical staining of Arabidopsis thaliana secondary cell wall elements. Journal of visualized experiments: JoVE Advance Access published May 13, 2014, doi:10.3791/51381.

Professional Counting Solutions for Your Industryhttps://countthings.com/en/ Accessed February 11, 2026.

Prusty, M. R., Bdolach, E., Yamamoto, E., Tiwari, L. D., Silberman, R., Doron-Faigenbaum, A., Neyhart, J. L., Bonfil, D., Kashkush, K., Pillen, K., et al. (2021). Genetic loci mediating circadian clock output plasticity and crop productivity under barley domestication. New Phytologist 230:1787–1801.

Rasmusson, D. C., and Wilcoxson, R. W. (1979). Registration of Morex Barley^1^ (Reg. No. 158). Crop Science 19:293–293.

Shalit-Kaneh, A., Eviatar-Ribak, T., Horev, G., Suss, N., Aloni, R., Eshed, Y., and Lifschitz, E. (2018). The flowering hormone Florigen accelerates secondary cell wall biogenesis to harmonize vascular maturation with reproductive development. bioRxiv Advance Access published 2018, doi:10.1101/476028.

Shalit-Kaneh, A., Eviatar-Ribak, T., Horev, G., Suss, N., Aloni, R., Eshed, Y., and Lifschitz, E. (2019). The flowering hormone florigen accelerates secondary cell wall biogenesis to harmonize vascular maturation with reproductive development. Proceedings of the National Academy of Sciences of the United States of America 116:16127–16136.

Tang, D., Chen, M., Huang, X., Zhang, G., Zeng, L., Zhang, G., Wu, S., and Wang, Y. (2023). SRplot: A free online platform for data visualization and graphing. PLOS ONE 18:e0294236.

Tao, W., Bian, J., Tang, M., Zeng, Y., Luo, R., Ke, Q., Li, T., Li, Y., and Cui, L. (2022). Genomic insights into positive selection during barley domestication. BMC Plant Biology 2022 22:1 22:1–19.

Tietze, H., Abdelhakim, L., Pleskačová, B., Kurtz-Sohn, A., Fridman, E., Nikoloski, Z., and Panzarová, K. (2025). Prediction of harvest-related traits in barley using high-throughput phenotyping data and machine learning. Front Plant Sci 16:1686506.

Waddington, C. H. (1942). Canalization of development and the inheritance of acquired characters. Nature 150:563–565.

Wang, Y., Lu, Y., Guo, Z., Ding, Y., and Ding, C. (2020). RICE CENTRORADIALIS 1, a TFL1-like gene, responses to drought stress and regulates rice flowering transition. Rice 13:1–12.

Zamir, D. (2001). Improving plant breeding with exotic genetic libraries. Nat Rev Genet 2:983–989.

